# BrainImageR: Spatiotemporal gene set analysis referencing the human brain

**DOI:** 10.1101/229302

**Authors:** Sara B. Linker, Jonathan Hsu, Adela Pfaff, Debha Amatya, Shu-Meng Ko, Sarah Voter, Quinn Wong, Fred H. Gage

## Abstract

**RATIONALE:** Neurological molecular analyses such as transcriptomics, epigenetics, and genome-wide association studies must be assessed in the context of the human brain in order to generate biologically meaningful inferences. It is often difficult to access primary human brain tissue; therefore, approximations are made using *in vitro* modeling or by identifying disease-associated genes from DNA extracted from blood. Gene sets from these studies are then compared to the post-mortem human brain to provide an assessment of the brain region and the developmental time point that a gene set is most closely associated with. However, most analyses of postmortem datasets are achieved by building new computational tools each time in-house, which can cause discrepancies from study to study, indicating that the field is in need of a user-friendly suite of tools to examine spatiotemporal expression with respect to the postmortem brain. Such a tool will be of use to the molecular interrogation of neurological and psychiatric disorders, with direct advantages for the disease-modeling and human genetics communities.

**RESULTS:** We have developed brainImageR, an R package that calculates both the spatial and temporal association of a dataset with post-mortem gene expression data from the Allen Brain Atlas. BrainImageR performs a robust analysis of gene set enrichment to identify enriched anatomical regions, provides a global high-resolution visualization of these enrichments across the human brain, and predicts when in developmental time the sample is most closely matched to, a task that has become increasingly important in the field of *in vitro* neuronal modeling. These functionalities of brainImageR enable a quick and efficient characterization of a given dataset across normal human brain development.

**AVAILABILITY AND IMPLEMENTATION:** BrainImageR is released under the Creative Commons CC BY-SA 4.0 license and the source code can be downloaded through github at https://github.com/saralinker/brainImageR.

## INTRODUCTION

Many neurological studies do not use primary tissue for their analysis. Genome-wide association studies are taken from the whole genome, which is largely independent of the tissue of origin, and transcriptomics and epigenetics studies are increasingly being assayed using *in vitro*-derived neurons. Therefore, it is important to have a clear and reproducible way to assess the relationship of this information to the primary tissue of the human brain. A common method is to compare genetic datasets to the post-mortem human brain using the Allen Brain Atlas reference data (Miller et al., 2014, Hawrylycz et al., 2012). Some packages and on-line tools are available to analyze this information (Grote et al., 2016, Stein et al., 2014), but these tools either lack the ability to predict developmental time, do not provide informative visualizations of the dataset with respect to the human brain, or require submitting data through an online portal that inhibits exploratory analysis and limits overall use. Therefore, researchers have turned to repeatedly building new computational tools in-house to perform spatiotemporal enrichment analyses (Lathe and Haas, 2017, Pasca et al., 2015, Pollen et al., 2014). This repeated effort indicates that the field requires an analysis suite that is easy to implement and is flexible across data types to identify the spatiotemporal relationship to the human brain. BrainImageR provides this flexible and intuitive resource within the R environment.

## SOFTWARE DESCRIPTION

### Spatial gene set enrichment

The Allen Brain Atlas assays gene expression across multiple regions of the brain throughout human development, ranging from 8 post-conception weeks (pcw) to 48 years of age (Miller et al., 2014, Hawrylycz et al., 2012). BrainImageR calculates the overlap between two lists, a user-supplied query and a post-mortem microarray reference from the Allen Brain Atlas (H376_IIIA_02, H0351.2001, H0351.2002, H0351.1009, H0351.1012, H0351.1015, and H0351.106) (Miller et al., 2014, Hawrylycz et al., 2012) (Figure 1A). The Allen Brain Atlas reference set notes the presence or absence of microarray probe signal for each gene. A gene is defined as expressed within a given tissue if at least 80% of the probes are active. The brainImageR function, *SpatialEnrichment,* takes as input the choice of using the developing or adult brain as well as a query gene list. *SpatialEnrichment* calculates the number of query genes that are expressed above baseline within each tissue and compares this value to either the entire background gene set or a user-defined background list. A bootstrapping procedure calculates the significance of enrichment over background. At this point the user will have an enrichment value and the associated bootstrapped p-value for all brain regions. The user can search this list directly for high-resolution information regarding the enriched tissue. For example, as a proof-of-concept, we curated a list of genes that are over-represented in either the ventral thalamus of the developing brain (Figure 1B) or the hippocampus of the adult brain (Figure 1C). BrainImageR returned the regional enrichment for every microdissected region along with the bootstrapped significance estimate, which for these gene sets was significant (p_adj_< 0.05) within the relevant thalamus and hippocampal brain regions.

**Figure 1.**
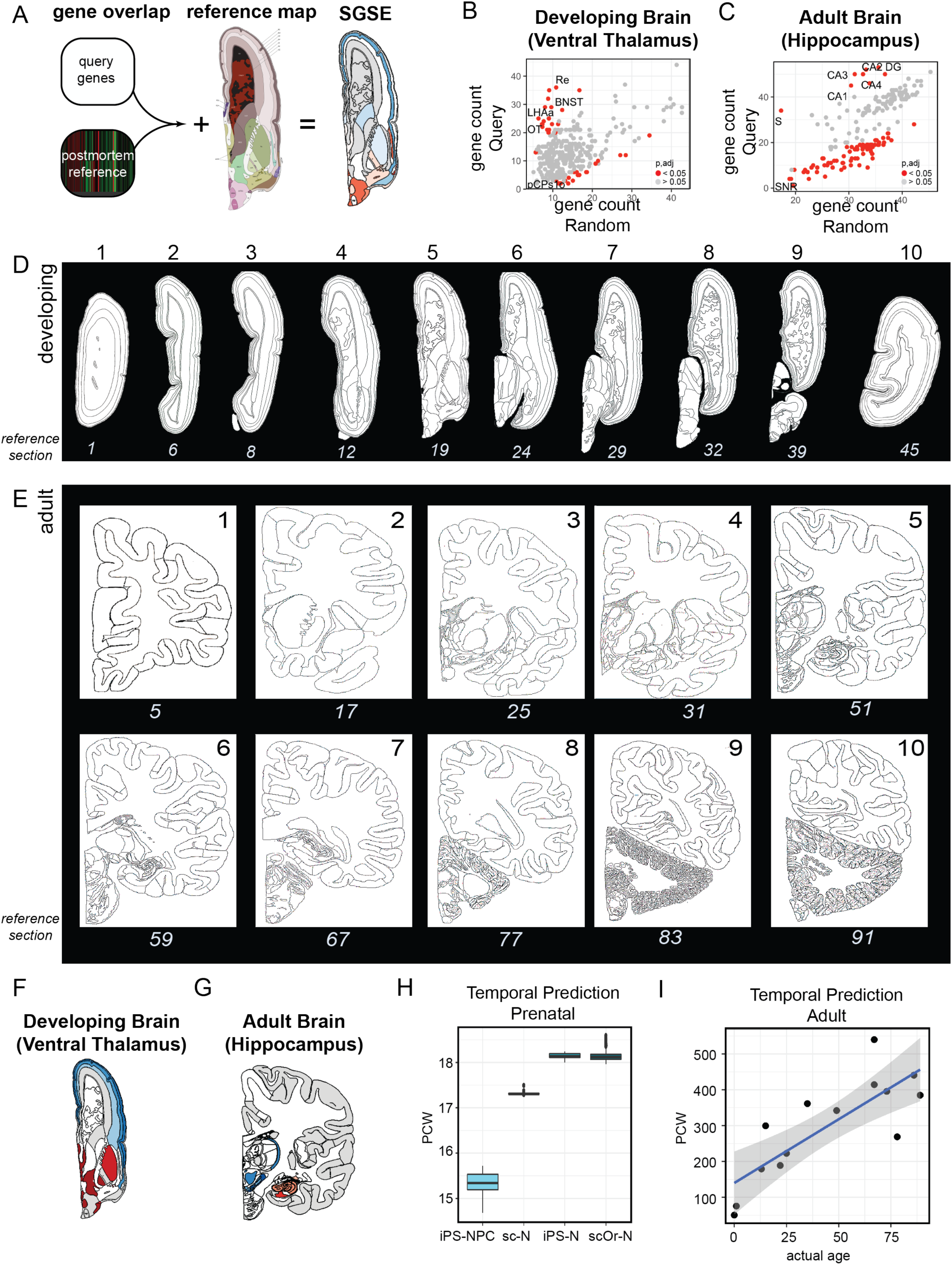
Spatial gene set enrichment with brainImageR. A) BrainImageR overlaps a query gene list with postmortem expression data from the Allen Brain Atlas. It then maps the enrichment values onto a reference image of the human brain. B-C) Example enrichment scores for each region when comparing the background gene set (x-axis) and the query gene set (y-axis) for the ventral thalamus of the developing brain (B) and the hippocampus of the adult brain (C). Red dots indicate enrichment scores with a bootstrapped p_adj_ < 0.05. D-E) Available reference images for the developing (D) and adult (E) brain. The top number is used as input into brainImageR to plot the enrichment data over the respective image. The bottom number is the section for the respective image in the Allen Brain Atlas and can be used to look up additional information about each section. F and G) BrainImageR visualization for the enrichment information in B and C, respectively. H) Predicted age in pcw from iPSC-neural progenitors (iPS-NPC), single-cell RNA-seq from post-mortem 16-18 pcw neurons (sc-N), 2D iPSC-neurons (iPS-N) and single-cell RNA-seq from 3D organoid neurons (scOr-N). I) Predicted age in pcw from human post-mortem prefrontal cortex.

An advantage of BrainImageR over other methods is the ability to map this gene set enrichment information to the reference brain, thereby creating a visualization of spatial gene set enrichment (SGSE). This visualization is achieved by referencing a conversion between the reference brain map and each tissue assayed by microarray in the Allen Brain Atlas.*CreateBrain* will use this information to transform the regional enrichment values into a brain map. The user can specify which section along the rostral-caudal axis they would like to plot within either the developing (Figure 1D) or adult (Figure 1E) brain. The user plots this composite graph as a tiff image with *PlotBrain*. For example, genes enriched in the ventral thalamus of the developing brain or the hippocampus of the adult show dark red for the respective regions (Figure 1F-G), indicating gene enrichment in these areas.

### Predicting developmental time given gene expression data

Developmental time-point prediction is particularly useful for *in vitro* modeling of neurons using induced-pluripotent stems cells both in 2D culture and 3D organoids. Importantly, when modeling neurons *in vitro*, it is difficult to identify what age in development the neurons represent. BrainImageR solves this problem by predicting the relative developmental time point by comparing the sample transcriptome information with the developmental transcriptome from the Allen Brain Atlas. For example, applying brainImageR to induced pluripotent stem cell (iPSC)-derived neuronal progenitor cells (Li et al., 2017) identifies a consistent prenatal signature with an increase in age as the neurons are differentiated either into neurons in the 2D culture setting (Schwartz et al., 2015) or into a 3D organoid setting assayed with single-cell RNA-sequencing (Camp et al., 2015) (Figure 1H). Furthermore, single-cell RNA-sequencing from post-mortem fetal cortex (16 - 18 pcw) was also predicted to be an average of 17.3 pcw (Darmanis et al., 2015) (Figure 1H), indicating the high accuracy of temporal prediction within the prenatal brain. The predict function can also be applied to adult post-mortem data, even though the temporal dynamics of the adult brain are limited compared to the developing brain. Although the accuracy is somewhat reduced in the adult predictions, the precision remains high and robust. For example, predicting developmental time from adult prefrontal cortex RNA-seq samples ranging from newborn to 89 years of age (Mertens et al., 2015) accurately identifies a linear association of predicted time with age (Figure 1I), indicating the robust precision of temporal estimates within the adult brain.

We have shown that brainImageR offers a suite of tools that provides the user with spatiotemporal enrichment information within the developing and adult human brain. The results are visualized in an easy-to-interpret anatomical map of the human brain. Further functionality of predicting the developmental time of each sample enables a complete characterization of *in vitro*-derived neurons with respect to the human post-mortem brain. With these tools, human genetics and neurological modeling studies will have a consistent framework within which to understand how a given dataset relates to gene expression within the human brain.

## ACKNOWLEDGEMENTS

We would like to thank Mary Lynn Gage for assistance in editing. Image credit: Allen Institute.

